# Two Septin Complexes Mediate Actin Dynamics During Cell Wound Repair

**DOI:** 10.1101/2023.11.14.567084

**Authors:** Viktor Stjepić, Mitsutoshi Nakamura, Justin Hui, Susan M. Parkhurst

## Abstract

Cells have robust wound repair systems to prevent further damage or infection and to quickly restore cell cortex integrity when exposed to mechanical and chemical stress. Actomyosin ring formation and contraction at the wound edge are major events during closure of the plasma membrane and underlying cytoskeleton during cell wound repair. Here, we show that all five *Drosophila* Septins are required for efficient cell wound repair. Based on their different recruitment patterns and knockdown/mutant phenotypes, two distinct Septin complexes, Sep1-Sep2-Pnut and Sep4-Sep5-Pnut, are assembled to regulate actin ring assembly, contraction, and remodeling during the repair process. Intriguingly, we find that these two Septin complexes have different F-actin bending activities. In addition, we find that Anillin regulates the recruitment of only one of two Septin complexes upon wounding. Our results demonstrate that two functionally distinct Septin complexes work side-by-side to discretely regulate actomyosin ring dynamics during cell wound repair.

## INTRODUCTION

As a result of physiological or environmental stresses, cells experience ruptures to their cortex (plasma membrane and underlying cortical cytoskeleton). To prevent further damage/death or infection, the cell must undergo a robust and efficient wound repair response to close the ruptured cortex and restore normal function ^1–11^. A major step after cell cortex injury is the formation and translocation of an actomyosin ring to pull the cortical cytoskeleton and overlying plasma membrane closed ^7,12,13^. The coordinated efforts of different actin filament populations, as well as their interactions with actin binding and regulatory proteins, are needed for proper spatiotemporal regulation of actomyosin ring construction and closure dynamics (cf. ^6^). One family of proteins that are known for their roles at the cell cortex and that associate with membranes of specific curvatures or lipid compositions, as well as with cytoskeleton networks of specific architectures or properties, is the Septin family ^14–23^.

Septins are a family of conserved GTP-binding proteins that form hetero-oligomeric complexes that polymerize into filaments (as well as into higher order structures including rings, gauzes, and collars) and that function in many cellular processes, including cytokinesis, epithelial wound healing, membrane remodeling, and when mis-regulated, are associated with neurodegenerative diseases and cancer ^16,18,19,21,24–27^. The number of Septin genes found in any organism can vary greatly from 13 in humans to 2 in *C. elegans* ^28,29^. Individual Septin family members are grouped into different classes (SEPT2, SEPT3, SEPT6, and SEPT7) based on sequence homology, with each class containing anywhere from 1 to 5 individual Septin family members ^30^. Septins form hetero-oligomeric complexes with each other and are often reported to function as tetramers (two dimers), hexamers (two trimers), or octamers (two tetramers) ^23,31–35^. The formed hetero-oligomeric complexes can in turn bind actin ^22,36,37^, microtubules ^22,38,39^, and lipid membranes ^40,41^. Septin-actin interactions have been investigated in budding yeast cytokinesis where Septins were found to be critical in the formation of actin cables across the bud neck ^42^ and stabilizing the cytokinetic ring ^43^.

*Drosophila* have 5 Septins that are grouped into three classes: Sep2 and Sep5 (SEPT6 class), Sep1 and Sep4 (SEPT2 class), and Pnut (SEPT7 class) (Fig. 1A) ^44^. As with Septins in other systems, *Drosophila* Septins have also been shown to bind actin and facilitate F-actin bending and actin ring compaction ^37,43^. *Drosophila* Septins have also been shown to modulate membrane topography and affect cell motility ^45^. In *Drosophila*, Septin complexes are reported as hetero-hexamers, where members of a single class are thought to be interchangeable (Fig. 1B) ^37,46,47^. However, recent evidence has shown that individual Septins have non-redundant functions during *Drosophila* oogenesis and *A. gossypii* development ^40,48^, suggesting that the various combinations of individual Septins within complexes may lead to distinct populations that behave differently and perform different functions ^47–49^.

**Fig 1.**
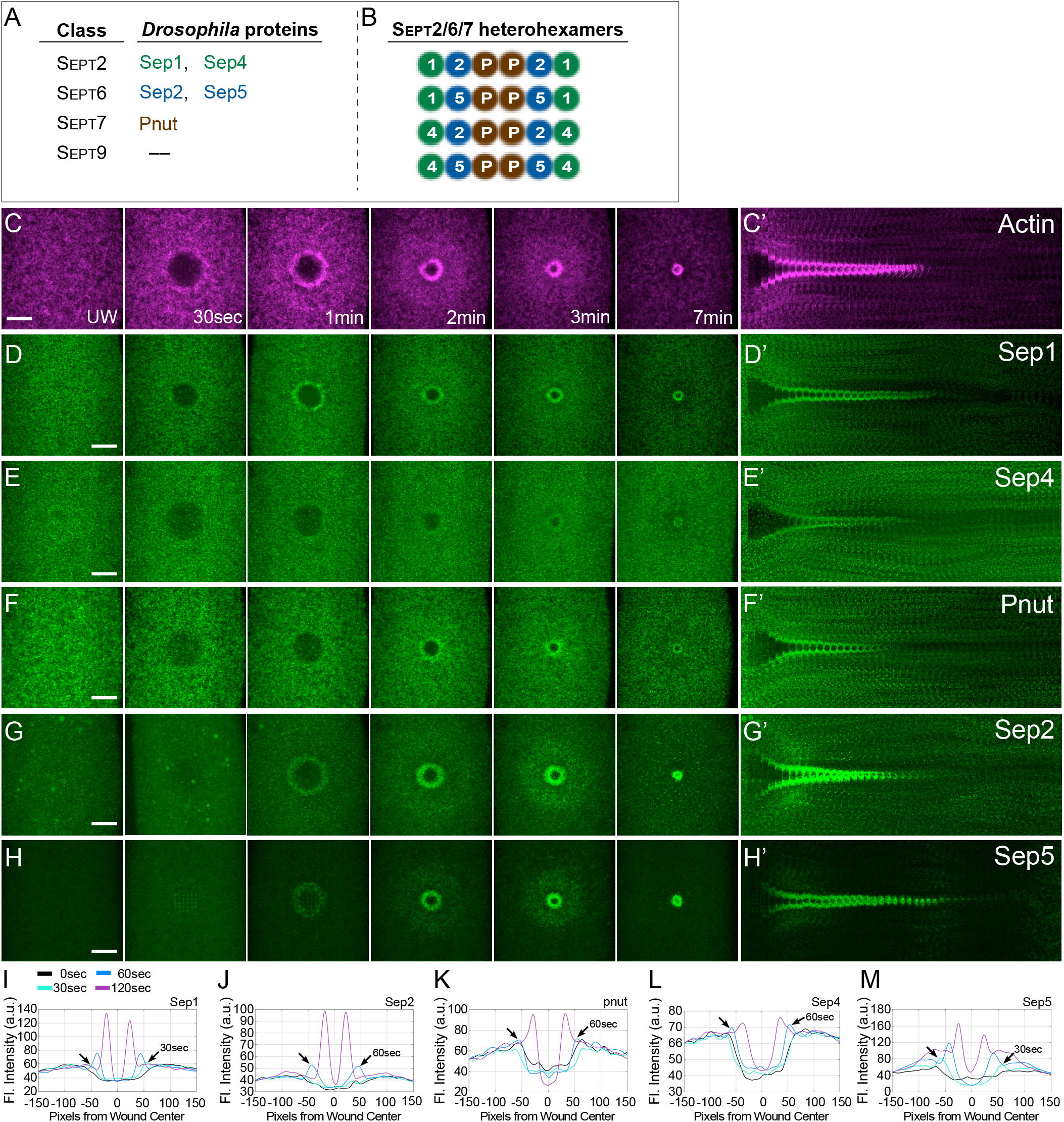
Individual Septins are recruited to the wound edge. (A) *Drosophila* Septins and their respective classes. (B) Potential *Drosophila* Septin hetero-hexamers. (C-H’) Confocal max projection images of embryos expressing an actin reporter (sGMCA or sStMCA; C-C’) and Sep1-GFP (D-D’), Sep4-GFP (E-E’), ChFP-Pnut (F-F’), Sep2-GFP (G-G’), or Sep5-RFP (H-H’), at the time points indicated. (C’-H’) Kymographs across the wound area in C-H, respectively. (I-M) Fluorescence intensity (arbitrary units) profiles across the wound area over time for the images shown in (C-H). Scale bars: 20μm.

Here we show that all 5 *Drosophila* Septins are recruited to wounds and are required for optimal cell wound repair in the *Drosophila* model. We find that individual knockdown of Septin family members results in impaired wound closure phenotypes that are not similar for members of the same class. Members of the same class also do not share similar recruitment patterns to the wound edge, suggesting non-redundant functions among members of the same Septin class. These distinct recruitment patterns and wound repair phenotypes suggest that two different Septin complexes (Sep1/Sep2/Pnut and Sep4/Sep5/Pnut) are necessary for optimal cell wound repair. Additionally, the Septin-interacting protein Anillin is required for cell wound repair, where it is needed for the proper recruitment of Sep1 and Sep2, but not Pnut, Sep4, or Sep5 to the wound edge. Lastly, we find that while individual Septins can bind to and bundle actin, Septin complexes are required to bend these bundled actin filaments into curved architectures. Taken together, our studies suggests that in addition to the potential roles for individual Septins, two distinct Septin complexes operate concomitantly for optimal cell wound repair.

## RESULTS

### Septins are recruited to cell wounds in spatially and temporally distinct patterns

To determine which members of the Septin family, if any, are recruited to the wound edge, we expressed either GFP- or mCherry-tagged Septin family members, along with an actin reporter (sGMCA or sStMCA; see Methods), in otherwise wildtype *Drosophila* syncytial embryos (Fig. 1C-N; Video 1; Table S1) ^50,51^. Upon laser injury on the lateral side of nuclear cycle 4-6 embryos (see Methods), actin begins to form an actomyosin ring at 30sec and by 1min a robust actomyosin ring is observed (Fig. 1C-C’; Video 1). Strikingly, upon laser injury all five *Drosophila* Septins are recruited to the wound edge in class-independent spatial patterns (Fig. 1D-M; Video1).

Interestingly, Sep1 and Sep2 are enriched in a ring at the wound edge that forms at the inner edge of the actomyosin ring and extend towards the center of the wound (Fig. 2A-B’, F-G). Sep4 and Sep5 are also enriched in a ring at the wound edge, but this ring overlaps completely with the actomyosin ring (Fig. 2D-E’, I-J). Pnut is recruited in a ring at the wound edge whose overlap with the actin ring is in between those observed for Sep1/Sep2 (internal to actin ring) and Sep4/Sep5 (overlapping actin ring) (Fig. 2C-C’, H). Thus, while all Septins are recruited to wounds, they exhibit distinct class-independent spatial recruitment patterns, suggesting that they likely play different roles in the cell wound repair process.

**Fig 2.**
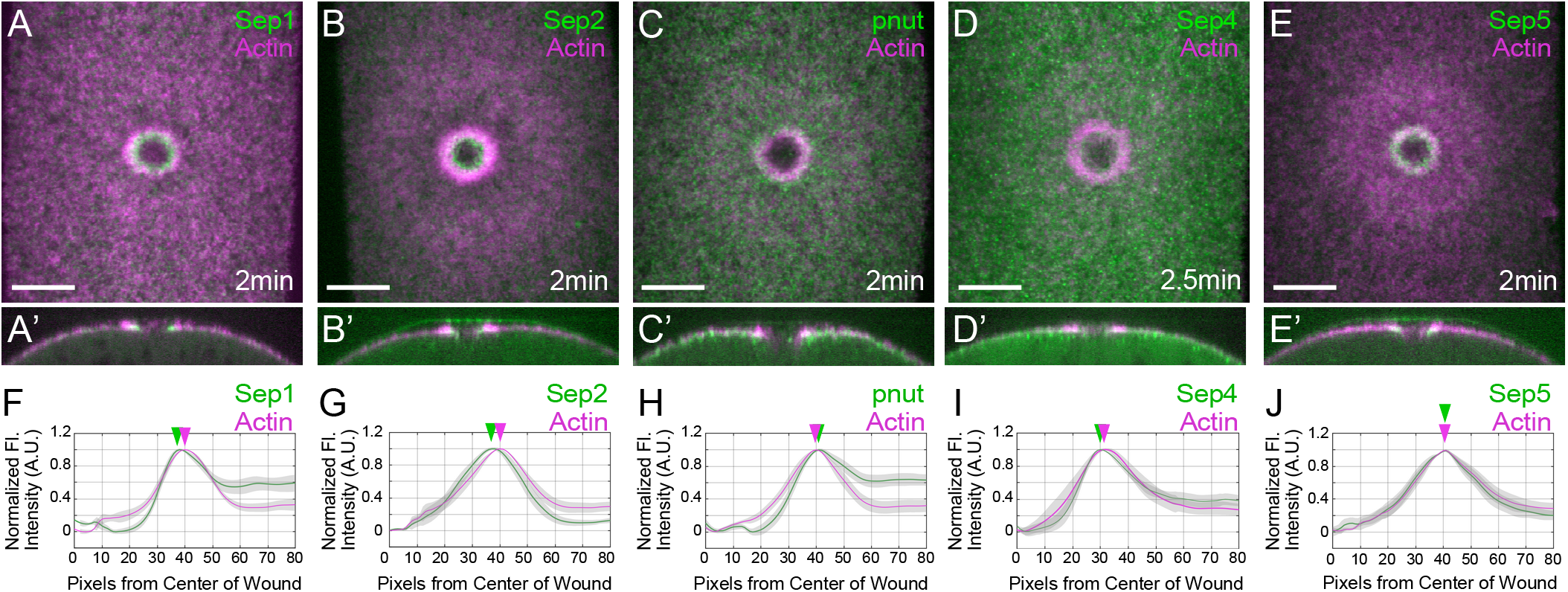
Spatial recruitment of individual Septins is class independent. (A-E) Confocal max projection images of embryos co-expressing an actin reporter (sGMCA or sStMCA) and Sep1-GFP (A), Sep2-GFP (B), ChFP-Pnut (C), Sep4-GFP (D), or Sep5-RFP (E). Scale bars: 20μm. (A’-E’) Orthogonal cross-sectional image of A-E, respectively. (F-J) Normalized fluorescence intensity (arbitrary units) average profiles (for n≥10) depicting localization of individual Septins with respect to the actin ring. Peak expression is indicated by the respective arrowheads.

### Knockdown of individual Septins results in impaired cell wound repair phenotypes

To assess the impact of each Septin on cell wound repair and actomyosin ring dynamics, we knocked down each Septin family member individually and followed actin dynamics by expressing an actin reporter (sGMCA) in the Septin knockdown backgrounds (Fig. 3; Tables S1, S2). We generated Septin knockdown embryos two different ways: 1) expressing RNAi constructs for Sep1, Sep4, or Pnut in the female germline using the GAL4-UAS system ^52,53^, and 2) using previously described null mutants for Sep2 and Sep5 (*Sep2*^2^ and *Sep5*^2^, respectively) ^54^ (see Methods). In control embryos, a robust actomyosin ring is formed by 1min post-wounding, which then contracts until the wound area is pulled closed ∼13min after injury and the actomyosin ring is then disassembled (Fig. 3A-A’, H, O-R; Video 2). Control embryos exhibit an initial 1.61±0.04-fold wound area expansion (Fig. 3O; Table S2) followed by a wound contraction rate of 7.23±0.30μm^2^/sec (Fig. 3P; Table S3). These embryos form robust actin rings (5.2±0.19μm in width) and with an actin ring intensity of 2.82±0.17 (Fig. 3Q-R; Table S2).

**Fig 3.**
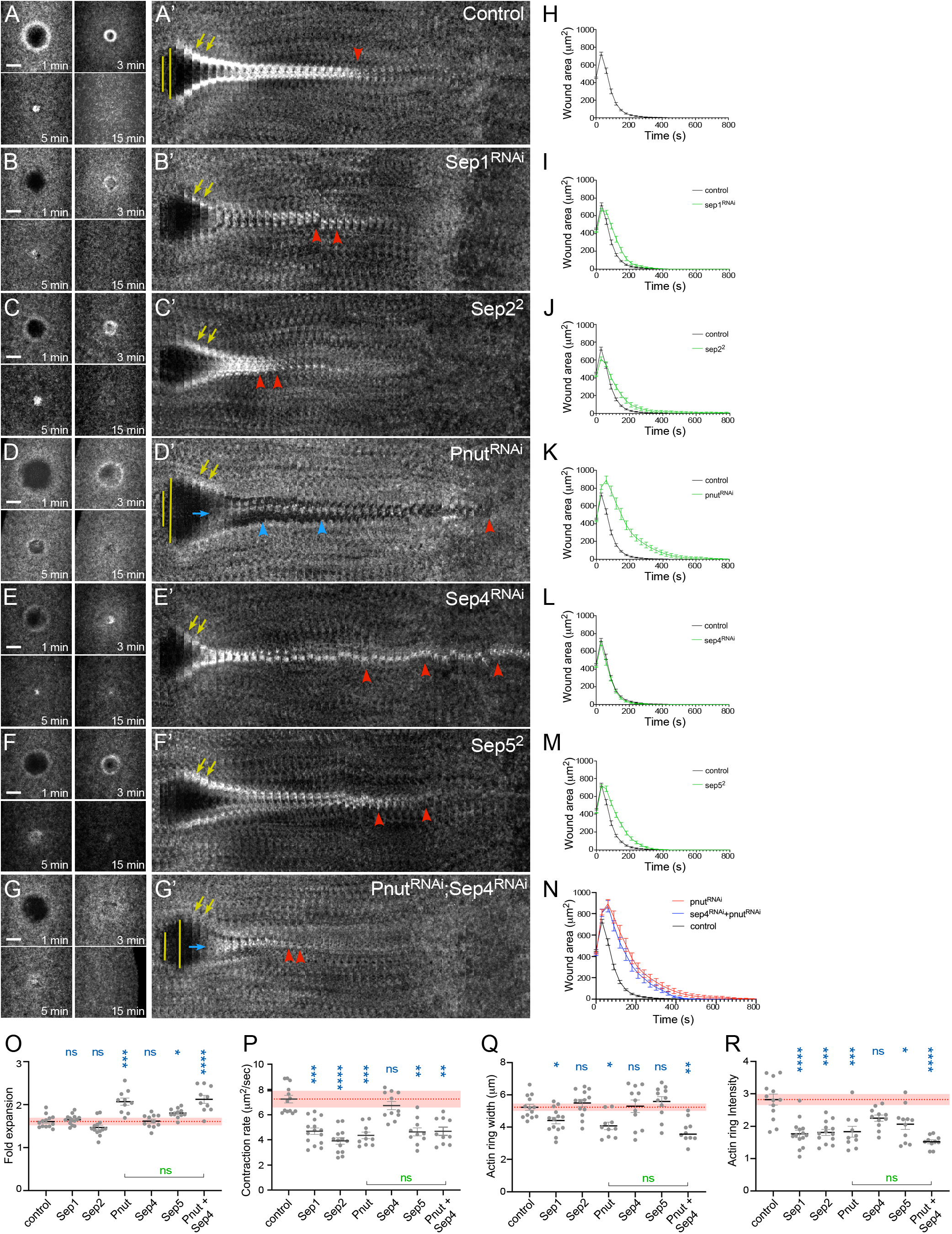
Knockdown of individual Septins result in class-independent wound repair phenotypes. (A-G’) Confocal max projection images of wounds generated in embryos expressing an actin marker (sGMCA) in control (vermilion RNAi; A), Sep1 RNAi (B), *Sep2*^2^ mutant (C), Pnut RNAi (D), Sep4 RNAi (E), *Sep5*^2^ mutant (F), and Pnut+Sep4 RNAi (G). (A’-G’) Kymographs across the wound area depicted in A-G, respectively. Wound expansion is highlighted by yellow lines. Actin recruitment to the actomyosin ring is indicated by yellow arrows. Actomyosin ring disassembly (or lack thereof) is indicated by red arrowheads. Actin accumulation internal to the wound is indicated by blue arrows. Failure of the wound to close is indicated by blue arrowheads. Scale bars: 20μm. (H-N) Quantification of the wound area over time for knockdowns shown in (A-G), respectively. (O-R) Quantification of actin ring dynamics in Septin knockdowns. Quantification of fold wound expansion (O), Wound contraction rate (P), actin ring width (Q), actin ring intensity (R). Black line and error bars represent mean ± SEM. Red dotted line and square represent mean ± 95% CI from control. Kruskal-Wallis test (blue) and Mann-Whitney test (green): * *p*<0.05, ** *p*<0.01, *** *p*<0.001, **** *p*<0.0001, ns is not significant. Comparison to control is indicated by blue asterisks; comparisons of individual pairs are indicated by a line and green asterisks.

Similar to their spatial recruitment patterns, knockdown of individual Septins resulted in different wound repair phenotypes (Fig. 3; Table S2; Video 2). Sep1 and Sep2 are both recruited to wounds overlapping the inside of the actomyosin ring and exhibit similar defects in actin dynamics during cell wound repair. Sep1 knockdown and *Sep2* mutant embryos exhibit delayed wound closure, reduced recruitment of actin to the actomyosin ring, and premature actomyosin ring disassembly (11min and 7min post wounding, respectively) compared to controls (13min post wounding) (Fig. 3A-C’, H-J, O-R; Table S2; Video 2). Similarly, Sep4 and Sep5 are both recruited to wounds overlapping the actomyosin ring and show similar defects in actin dynamics to each other, but different from those exhibited by Sep1 and Sep2. Sep4 knockdown and *Sep5* mutant embryos exhibit delayed actomyosin ring disassembly (>30min and 20min post wounding, respectively) compared to controls (13min post wounding) (Fig. 3E-F’, L-M, O-R; Table S2; Video 2). Pnut, which is also recruited in a ring mostly overlapping the actomyosin ring, had the most severe effect on actin ring dynamics when knocked down. Pnut knockdown resulted in wound over-expansion, reduced actin recruitment to the wound edge, actin recruitment to the center of the wound, and delayed wound closure (Fig. 3D-D’, K, O-R; Table S2; Video 2). While all five *Drosophila* Septins are required for optimal cell wound repair, Sep1/Sep2 have similar spatial recruitment patterns and exhibit similar effects on actin dynamics when knocked down. A similar result was observed for Sep4/Sep5, suggesting that there may be two major Septin complexes – Sep1/Sep2/Pnut and Sep4/Sep5/Pnut – functioning in the context of cell wound repair.

*Drosophila* Septin complexes are hexamers made of two trimers, that in turn consist of one member from each Septin class. In *Drosophila*, Pnut is the only member of the SEPT7 class (Fig. 1A-B) and may be vital for all Septin complex formation. If Pnut is indispensable for trimer and/or oligomeric Septin complex formation, knockdown of Pnut and one other Septin should exhibit a similar phenotype as Pnut alone. To investigate this possibility, we knocked down Pnut and Sep4 simultaneously and tracked actin ring dynamics. Double knockdown embryos exhibited wound repair defects similar to Pnut knockdown alone: wound over-expansion, reduced actin recruitment to the wound edge, actin recruitment to the center of the wound, reduced actin to the actomyosin ring, and delayed wound closure (Fig. 3G-G’, N-R; Table S2; Video 2). Taken together, our results suggest that Pnut is likely part of all Septin complexes in the context of cell wound repair in the *Drosophila* model.

### Two distinct Septin complexes are functioning concomitantly during cell wound repair

When one Septin within a Septin complex is knocked down, it has been shown to affect the expression of the other Septins in that complex, but not Septins in unrelated complexes ^34,35,46^. As our data suggests that two distinct Septin complexes—Sep1/Sep2/Pnut and Sep4/Sep5/Pnut—are likely functioning during cell wound repair, we analyzed the recruitment of Sep2, Sep5, and Pnut in the background of either Sep1 or Sep4 knockdowns (Fig. 4). In a Sep1 knockdown, we find that Sep5 and Pnut, but not Sep2, are recruited to the wound edge (Fig. 4A-F, J-L). Conversely, we find that in a Sep4 knockdown, Sep2 and Pnut, but not Sep5, are recruited to the wound edge (Fig 4G-L). Taken together, our findings further suggest that Sep1/Sep2/Pnut and Sep4/Sep5/Pnut are functioning as distinct complexes within the context of cell wound repair.

**Fig 4.**
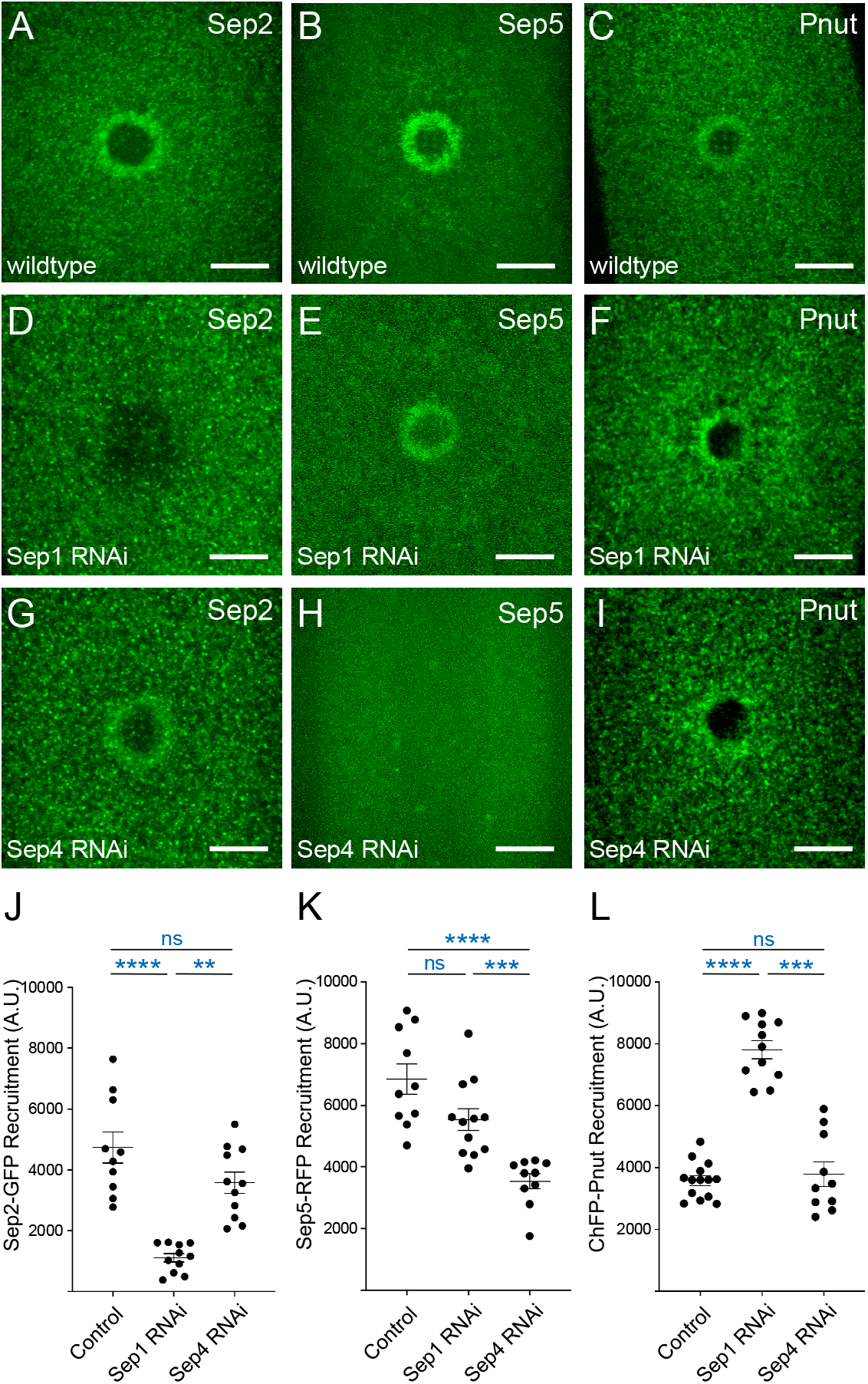
Two distinct Septin complexes are functioning during cell wound repair. (A-I) Confocal max projection images of wounds generated in embryos expressing Sep2-GFP (A, D, G), Sep5-RFP (B, E, H), or ChFP-Pnut (C, F, I) in wildtype (A-C), Sep1 RNAi knockdown (D-F), or Sep4 RNAi knockdown (G-I) backgrounds. Scale bar: 20 µm. (J-L) Septin recruitment (area under the curve of fluorescence profile) for Sep2-GFP (J), Sep4-GFP (K), or ChFP-Pnut (L) in control (wildtype), Sep1 RNAi, and Sep4 RNAi backgrounds at 70% wound closure (100% = fully open). Error bars represent ±SEM. Kruskal-Wallis test was performed for J-L. Comparison to control is indicated by blue asterisks: * *p*<0.05, ** *p*<0.01, *** *p*<0.001, **** *p*<0.0001, ns is not significant.

### Septin complexes crosslink actin filaments into curved bundles, whereas individual Septins bundle but do not bend actin filaments

Prior studies have established that both the *Drosophila* Sep1-Sep2-Pnut complex and the human Sep2-Sep6-Sep7 complex possess the ability to interact with F-actin, promoting actin bundling and generation of curved actin filament networks ^37,55,56^. Since both Sep1/Sep2/Pnut and Sep4/Sep5/Pnut complexes affect F-actin dynamics during cell wound repair, we examined the ability of these purified Septin complexes to generate curved actin bundles using actin polymerization assays (see Methods) (Fig. 5). When no purified proteins are added to these *in vitro* assays (no protein control), single actin filaments are randomly dispersed on the slide without forming bundles or higher order structures (Fig. 5A-B). Consistent with a previous study, we find that addition of purified Sep1/Sep2/Pnut protein complex resulted in actin filament bundling and bending to form circular structures (Fig. 5C-D). To quantify actin circle formation, we determined the average number of actin circles formed per field of view (Fig. 5S). Interestingly, we find that addition of purified Sep4/Sep5/Pnut protein complex also resulted in actin filament bundling and bending to form circular structures (Fig. 5E-F). No significant difference in the number of circles formed per field of view was observed between the two Septin complexes (Fig. 5S). Strikingly, we find a significant difference in the area of the circles formed by the two Septin complexes: 2.3±0.3 µm^2^ for Sep1/Sep2/Pnut complexes and 6.1±0.9 µm^2^ for Sep4/Sep5/Pnut complexes (p≤0.001) (Fig. 5T). These results indicate that the two Septin complexes are bending F-actin bundles to different degrees to achieve different curvatures.

**Fig 5.**
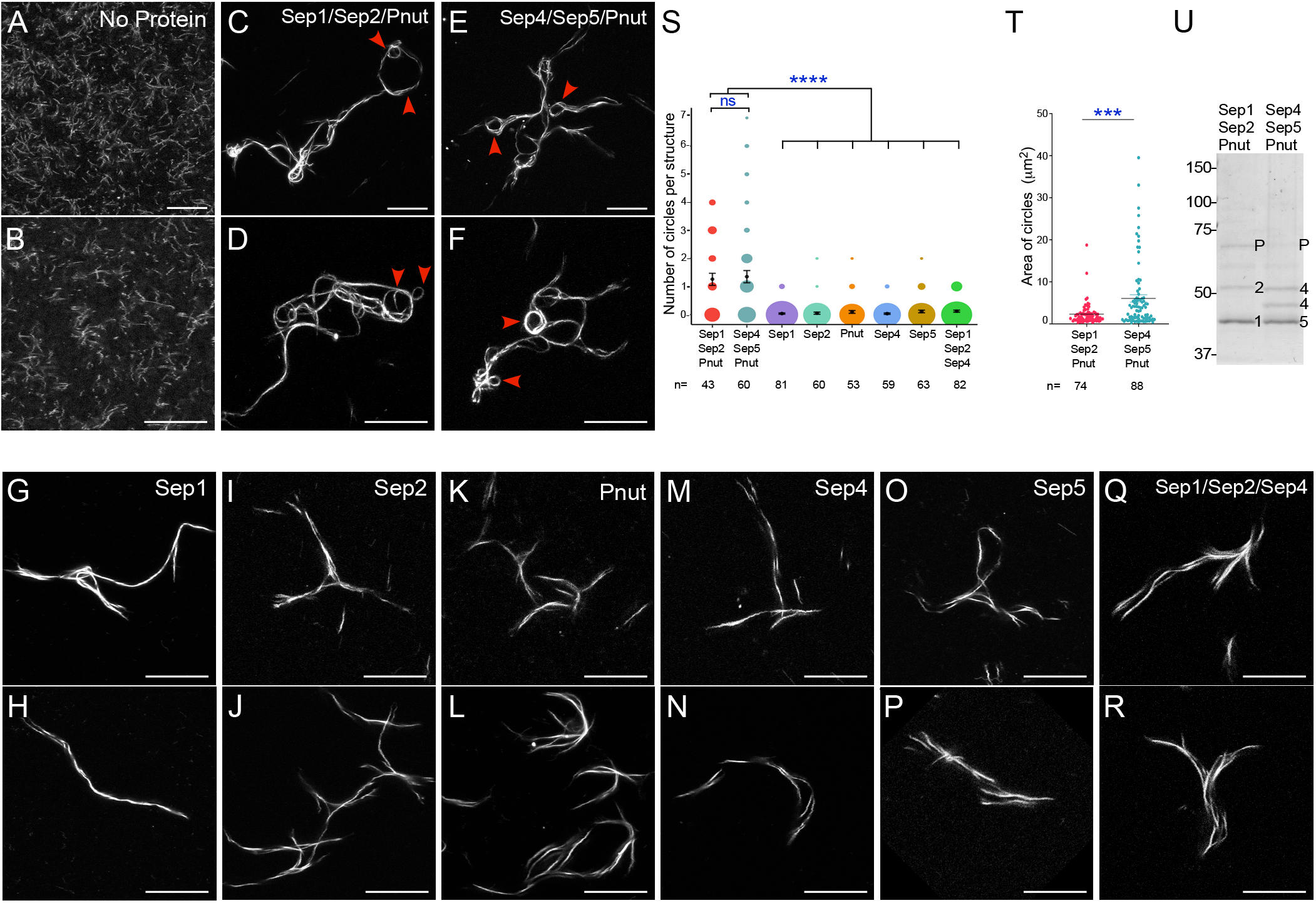
Septin complexes crosslink actin filaments into curved bundles, whereas individual Septins bundle but do not bend actin filaments. (A-R) *in vitro* generated actin filaments in the presence of no added protein (A-B), Sep1/Sep2/Pnut complex (C-D), Sep4/Sep5/Pnut complex (E-F), Sep1 alone (G-H), Sep2 alone (I-J), Pnut alone (K-L), Sep4 alone (M-N), Sep5 alone (O-P), or Sep1 + Sep2 + Sep4 proteins (Q-R). Circular structures formed in the presence of the Sep1/Sep2/Pnut or Sep4/Sep5/Pnut complexes are indicated by red arrowheads. (S) Quantification of actin circles per field of view formed by the individual or complexes of Septins as indicated. (T) Quantification of the area of the actin circles per field of view formed by each of the two Septin complexes as indicated. P value and number of structures assayed (n=) are indicated. (U) Coomassie stained gel of the purified Sep1/Sep2/Pnut and Sep4/Sep5/Pnut complexes. Septin proteins are indicated: 1=Sep1, 2=Sep2, 4=Sep4, 5=Sep5, P=Pnut. Scale bars: 20 µm. Error bars represent ±SEM. Kruskal-Wallis test was performed for S; Mann-Whitney test was performed for T. Comparison to control is indicated by asterisks: *** *p*<0.001, **** *p*<0.0001, ns is not significant.

We then examined the potential role that individual Septins may play in actin filament dynamics. Strikingly, we find that all five *Drosophila* Septins can bind to and bundle actin filaments, however they do not give rise to circular actin geometries (Fig. 5G-P, S). To determine if the circular actin filament geometries generated by the Septin complexes were due to functional complex formation or the additive effect of the three individual Septins present in each complex, we performed the same assay but with the addition of Sep1, Sep2, and Sep4 individually purified proteins. We find that the presence of three random purified Septin proteins does not result in actin filament bundling and curvature to form circular structures (Fig. 5Q-S), suggesting that the interaction of the three Septin proteins within the purified Septin complexes is synergistic rather than additive.

### Anillin is required for proper cell wound repair

Anillin binds to both actin and Septins and is thought to crosslink them ^57–65^. Anillin has been shown to promote the compaction of the diffuse Septin meshwork into well-organized rings at the actomyosin ring during cytokinesis ^43,61,64,66–69^. Therefore, we analyzed whether Anillin is recruited to cell wounds using embryos expressing a GFP-tagged version of Anillin along with an actin reporter (sStMCA; see Methods) (Fig. 6A-D; Video 3). Upon laser injury, Anillin forms a robust ring as early as 1min and closely follows actin ring dynamics (Fig. 6A-D; Video 3). This Anillin ring around the periphery of the wound overlaps with the inner edge of the actomyosin ring and extends into the center of the wound (Fig. 6C-D; Video 3). This is consistent with Anillin being a downstream effector of Rho1 which exhibits a similar recruitment pattern ^61,70^. Unexpectedly, Anillin is also recruited to the actin halo region which is where Cdc42 and Rac1 have been shown to be recruited ^70^ (Fig. 6A-D), suggesting that Anillin may also be interacting with Cdc42 and/or Rac1, as well as Rho1. Therefore, we wounded embryos expressing GFP fused to the Rho binding domain (RBD) of Anillin along with an actin reporter (sStMCA) (Fig. S1A-C). Consistent with the recruitment pattern of GFP-tagged full length Anillin, the Anillin RBD is also recruited to the inner edge of the actomyosin ring as well as the actin halo region. Performing a GST pulldown with *in vitro* translated Anillin RBD and GST-fused Rho1, Rac1, and Cdc42 showed that the Anillin RBD is not specific to GTP-bound Rho1 but binds to both GDP- and GTP-bound Rho1, Cdc42, and to a lesser extent Rac1 (Fig. S1D-E).

**Fig 6.**
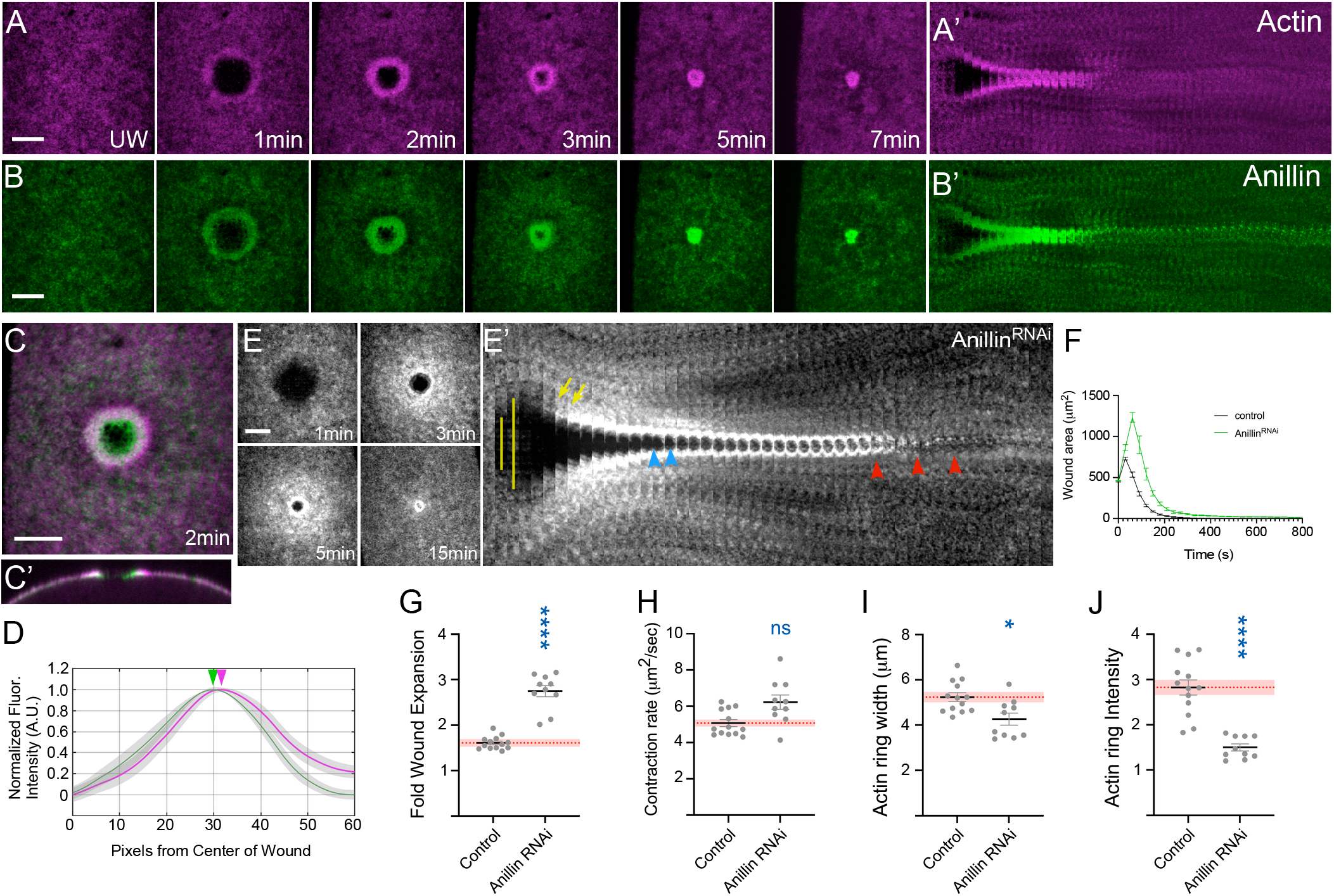
Anillin is required for optimal wound repair. (A-C’) Confocal max intensity projection images of embryos expressing an actin marker (sStMCA; A), GFP-Anillin (B), or both (C). (A’-B’) Kymographs across the wound area in A-B, respectively. (C’) Orthogonal cross-sectional image of an embryo expressing sStMCA and GFP-Anillin. (D) Fluorescence intensity (arbitrary units) average profile (for n=10) depicting recruitment of Anillin with respect to the actin ring. Peak expression is indicated by the respective arrowheads. (E) Confocal max projection images of wound generated in embryos expressing an actin marker (sGMCA) in an Anillin RNAi background. (E’) Kymograph across the wound area depicted in E. Wound expansion is highlighted by yellow lines. Actin recruitment to the actomyosin ring is indicated by yellow arrows. Delayed actomyosin ring disassembly is indicated by red arrowheads. Delay in wound closure is indicated by blue arrowheads. Scale bars: 20μm. (F) Quantification of the wound area over time for knockdown shown in (E). (G-J) Quantification of fold wound expansion (G), wound contraction rate (H), actin ring width (I), and actin ring intensity (J). Black line and error bars represent mean ± SEM. Red dotted line and square represent mean ± 95% CI from control. Kruskal-Wallis test: * *p*<0.05, ** *p*<0.01, *** *p*<0.001, **** *p*<0.0001, ns is not significant. Scale bars: 20 µm.

To investigate how Anillin affects actin ring dynamics, we examined wounds in an Anillin RNAi knockdown background and tracked actin dynamics using a fluorescent actin reporter (sGMCA). Consistent with its recruitment, Anillin knockdown embryos exhibited significant impairment in their ability to undergo repair (Fig. 6E-J; Table S2; Video 3). Notably, cells lacking Anillin exhibited a significantly greater initial fold wound expansion and lower actin ring intensities (Fig. 6G-J; Table S2). In contrast, minimal changes were observed in both the contraction rate and actin ring width (Fig. 6H-I; Table S2). The less intense actin ring is consistent with previous studies in the cytokinetic ring demonstrating that Anillin is also necessary for actomyosin ring compaction during cell wound repair.

### Anillin functions upstream of Sep1 and Sep2, but not Sep4, Sep5, or Pnut

Previous studies have suggested that Septin recruitment depends on Anillin ^62,64,67,71,72^. To distinguish among the roles of different Septins and/or Septin complexes, we examined individual Septin recruitment in the absence of Anillin and vice versa. Embryos co-expressing GFP-tagged Anillin and an actin marker were wounded in the background of individual Septin knockdowns. As expected, Anillin recruitment to wounds is not affected by the individual knockdown of Sep1, Pnut, and Sep4 (Fig. 7A-D). In contrast, differences are observed in the recruitment of individual Septins in an Anillin knockdown background (Fig. 7E-J). Sep1 and Sep2 are no longer recruited to the wound edge without Anillin, whereas Sep 4, Sep5, and Pnut did not exhibit a significant change in their recruitment (Fig. 7E-J). Combined with the similar spatial localizations of Sep1, Sep2, and Anillin at the inner edge of actin ring, our studies suggest an important role for Anillin in facilitating the recruitment of the Sep1/Sep2/Pnut complex, but not the Sep4/Sep5/Pnut complex, to the wound edge.

**Fig 7.**
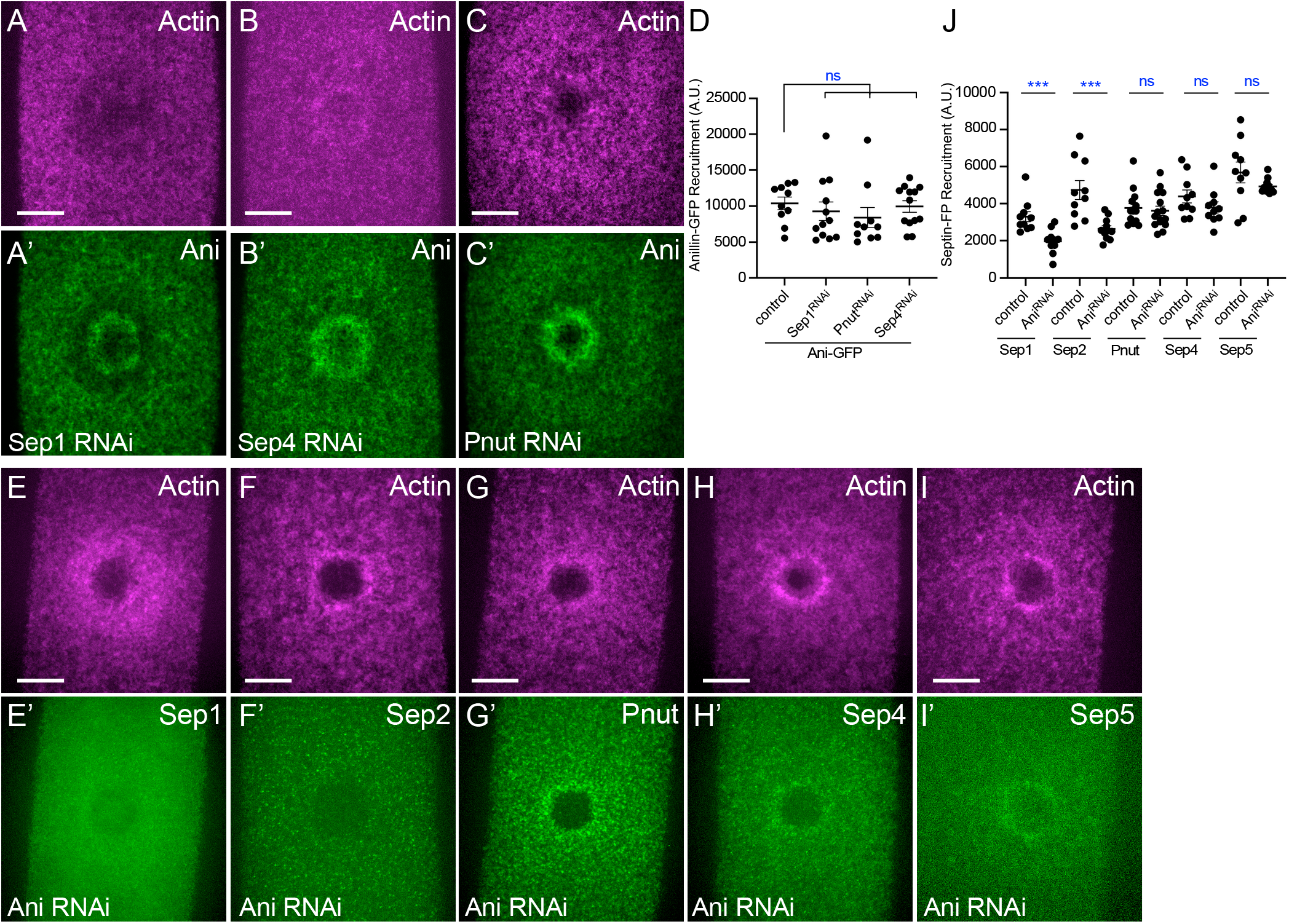
Anillin functions upstream of Sep1 and Sep2, but not Sep4, Sep5, or Pnut. (A-C) Confocal max projection images of wounds generated in embryos expressing an actin reporter (A-C) and GFP-Anillin (A’-C’) in Sep1 RNAi knockdown (A-A’), Sep4 RNAi knockdown (B-B’), or Pnut RNAi knockdown (C-C’) backgrounds. (D) Quantification of GFP-Anillin recruitment (area under the curve of fluorescence profile) in control, Sep1 RNAi, Sep4 RNAi, and Pnut RNAi backgrounds at 50% wound closure. (E-I) Confocal max projection images of wounds generated in embryos expressing an actin reporter (E-I) and Sep1-GFP (E’), Sep2-GFP (F’), ChFP-Pnut (G’), Sep4-GFP (H’), or Sep5-RFP (I’) in an Anillin RNAi knockdown background. (J) Quantification of Sep1-GFP, Sep2-GFP, ChFP-Pnut, Sep4-GFP, and Sep5-RFP recruitment (fluorescence intensity) in control and Anillin RNAi backgrounds at 50% wound closure. Error bars represent ±SEM. Kruskal-Wallis test was performed for D and J. Comparison to control is indicated by blue asterisks: * *p*<0.05, ** *p*<0.01, *** *p*<0.001, **** *p*<0.0001, ns is not significant. Scale bar: 20 µm.

## DISCUSSION

Cell wound repair in various organisms and systems has demonstrated a conserved requirement for the formation and translocation of an actin ring to achieve wound closure. Previous studies have demonstrated an important role for actin organization and architecture in the contractility of actomyosin rings ^6,73,74^. We find that Septin family proteins are required for optimal cell wound repair. While many studies have focused on the role of a specific Septin or Septin complex in a particular context, we show that all Septin family members are recruited to the cell wound edge, and are required for actin ring formation, wound closure, and remodeling during cell wound repair. Based on their recruitment patterns to wounds and wound repair phenotypes in knockdowns/mutants, we find that two Septin complexes (Sep1/Sep2/Pnut and Sep4/Sep5/Pnut) function concomitantly in different regions around the wound. In addition to the previous finding of F-actin bundling and bending activity in Sep1/Sep2/Pnut complex ^37^, our *in vitro* experiments show similar activity in Sep4/Sep5/Pnut complex. Intriguingly, these two Septin complexes bend F-actin to different degrees. Finally, we show that Anillin is upstream of the Sep1/Sep2/Pnut complex, but not the Sep4/Sep5/Pnut complex. Our results suggest that two distinct pathways regulate Septin recruitment to the wound in order to form two separate Septin complexes regulating different aspects of actin dynamics such as assembly, contraction, and remodeling during cell wound repair.

Septins are subdivided into four classes and assemble into hetero-oligomers containing one member from each of its different classes. These hetero-oligomers can then assemble into high-order structures such as filaments, gauzes, and rings to regulate cytoskeleton dynamics ^21,23,26,35^. Cryo-EM studies of the human Septin hetero-hexamer complex have shown that it tends to bend at its center, which is thought to induce curvature of membranes to which it is attached ^75^. Interestingly, we observe curved F-actin bundles induced by two separate Septin complexes, whereas individual Septins can only bind to and bundle F-actin. Thus, the bent conformation of Septin hexamers likely induced the observed circular actin bundles, in stark contrast to actin bundled by individual Septins. Intriguingly, we show that the diameter of actin circles generated by the Sep4-Sep5-Pnut complex is greater than those generated by the Sep1-Sep2-Pnut complex, indicating that the two Septin complexes have different propensities to curve F-actin. Depending on their position within the hetero-hexamers, Sep1 and Sep4 might exhibit different degrees of bending that lead to different actin filament curvatures. Alternatively, outer sides that contain Sep2-Pnut or Sep5-Pnut complexes might have different conformational flexibilities that limit or enhance the curvature when they form high-order structures. Further cryo-EM studies are needed to understand how the different Septin complexes bend F-actin, as well as how individual Septins affect this bending activity and the formation of high-order structures.

In addition to different propensities to bend F-actin, our results indicate that the two Septin complexes have distinct roles during cell wound repair. Sep1 and Sep2 are recruited inside the actin ring. Rho1 GTPase and the formin Diaphanous are also recruited inside the actin ring where they polymerize linear actin at the wound edge ^6,70,76^. As we showed previously that F-actin architecture is severely altered in *Drosophila* Arp2/3 knockdown backgrounds ^6^, actin is not simply polymerized along with the wound edge to form the actin ring. Rather actin needs to be bent in order to be integrated into the actin ring. Since the inside edge of the actin ring exhibits higher curvature compared with that of the outside edge of the ring, the higher curvature activity of the Sep1-Sep2-Pnut complex may be required for actin ring assembly. In addition, the specific F-actin bundling activity of each Septin is required for proper actin dynamics during cell wound repair. We find that actin ring organization is more disrupted in Sep1 and Sep2 knockdown/mutant backgrounds than those of Sep4 and Sep5. Septin complexes may also contribute to generating the necessary force required for wound closure. While contraction by actin and myosin is a major source of force for the actin ring closure, the bending of F-actin within the actin ring may alter the conformation of actin filaments. Such changes to their physical properties could alter the amount of force or the exact mechanisms of force generation that can be or need to be applied by force generating molecules. Once the actomyosin ring has contracted and closed the wound, it needs to be disassembled to restore the homeostatic state. Interestingly, Sep4 and Sep5 knockdown/mutant exhibit delayed actin ring remodeling implicating their roles in actin disassembly. The Sep4-Sep5-Pnut complex overlapping with the actin ring might give more flexibility to the ring by bending filaments to allow more efficient binding of disassembly factors to enhance disassembly (i.e., cofilin severs bent actin filament regions more efficiently)^77,78^.

Anillin has actin and Septin binding sites and functions as a scaffold protein to regulate actin dynamics during cytokinesis. Anillin has been shown to play a critical role in stabilizing and organizing diverse Septin architectures within the cytokinetic ring and during neuron development ^43,79^. Studies from fission yeast have shown that while Septins initially manifest as a diffuse network, flanking the contractile ring, Anillin orchestrates the compaction of Septins. This compaction allows the necessary rigidity and compact organization within the interior of the actin ring, thereby facilitating effective ring contraction and ensuring membrane rigidity ^43,67^. The absence of this compaction disrupts cytokinesis, allowing the diffuse Septin meshwork to anomalously persist into the subsequent cycle ^43^. Consistent with these findings, Anillin is also recruited to the inner edge of the actin ring during cell wound repair suggesting its similar role in Septin compaction and actin ring dynamics. Additionally, we find that Anillin is required for cell wound repair and the recruitment of Sep1 and Sep2 to the inside of the wound. While previous studies showed that Anillin knockdowns disrupt Sep2 and Pnut recruitment to the equator during cytokinesis, we find that Sep4, Sep5, and Pnut are still recruited to cell wounds. These results indicate that Septin recruitment to the wounds is regulated by at least two distinct pathways and Anillin is only required for the function of the Sep1-Sep2-Pnut complex inside of the actin ring. Since Anillin is still recruited to the wound in Sep1 and Sep2 knockdown/mutant backgrounds, Anillin might work as a scaffold to assemble Sep1 and Sep2 into the Sep1-Sep2-Pnut complex and/or compact the network of these proteins at the wound periphery. Identification of other proteins containing actin and Septin binding sites is needed to fully understand the roles of the Sep4-Sep5-Pnut complex pathway during cell wound repair.

Intriguingly, the wound repair phenotypes observed in Anillin knockdown embryos differ from Sep1 and Sep2 knockdowns even though our results show that Anillin is upstream of Sep1 and Sep2. These results suggest that Sep1 and Sep2 are not the only downstream effectors of Anillin and that Anillin is likely performing multiple functions during cell wound repair. Consistent with this, we show that the Rho binding site in Anillin can also bind Rac and Cdc42. Additionally, the Anillin recruitment pattern overlaps with those of Rho1, Rac1, and Cdc42 ^80^. Thus, Anillin might work with other GTPases during cell wound repair, which may result in wound expansion and faster wound contraction phenotypes in Anillin knockdowns.

Overall, we find that two distinct Septin complexes concomitantly regulate different aspects of actin dynamics during cell wound repair and have different F-actin bending activities. The intricate interplay of Septins is also observed in synaptic dysfunction leading to neurodegenerative and neurobehavioral disorders ^81^. Super-resolution microscopy, cryo-EM/-ET, biochemistry, and genetic approaches are needed to address remaining questions such as how the F-actin bending activity of Septin complexes contributes to actin ring assembly, contraction, and remodeling, and what molecules regulate the Sep4-Sep5-Pnut complex during cell wound repair.

## ACKNOWLEDGEMENTS

We thank Jihong Bai, Toshio Tsukiyama, Tony Cooke, Gerry Smith, Denise Clarke, Andrew Wilde, Thomas Lecuit, Tessa Allen, and Parkhurst lab members, the Bloomington Stock Center, the Harvard Transgenic RNAi Project, FlyBase, the Developmental Studies Hybridoma Bank, and the Fred Hutch/Leica Center of Excellence for advice, microscopes, antibodies, DNAs, flies, and other reagents used in this study. This work was supported by NIH grant GM111635 and the Mark Groudine Chair for Outstanding Achievements in Science and Service (to SMP) and, in part, through the NCI Cancer Center Support Grant P30 CA015704 (Shared Resources).

The authors declare no competing financial interests.

## Author Contributions

All authors contributed to the design of the experiments, performed experiments, analyzed data, and contributed to the writing of the manuscript.

## MATERIALS AND METHODS

**Reagents used in this study are described in Table S1.**

### Fly stocks and genetics

Flies were cultured and crossed at 25°C on yeast-cornmeal-molasses-malt extract medium. Flies used in this study are listed in Table S1. RNAi lines were driven using the maternally expressed GAL4-UAS driver, Pmatalpha-GAL-VP16V37.

An actin reporter, sGMCA (spaghetti squash driven, moesin-alpha-helical-coiled and actin binding site fused to GFP) reporter ^50^. or the mScarlet-i fluorescent equivalent, sStMCA ^51^, was used to follow wound repair dynamics of the cortical cytoskeleton.

In this study, we used an actin reporter + maternal GAL4 driver + *vermilion* RNAi (unrelated fly RNAi) as the control. RNAi knockdowns were quantified by qPCR (including biological and technical replicates).

Localization patterns and mutant analyses were performed at least twice with independent genetic crosses and ≥10 embryos were examined. Images representing the average phenotype were selected for figures.

### Quantification of RNAi knockdown eficiencies

RNAi knockdown efficiency was quantified using either qPCR. To harvest total RNA for qPCR, 100–150 embryos were collected after a 30 min incubation at 25°C, treated with TRIzol (Invitrogen/Thermo Fisher Scientific) and then with DNase I (Sigma). 1 μg of total RNA was reverse transcribed using the iScript gDNA Clear cDNA Synthesis Kit (Bio-Rad). RT-PCR was performed using the iTaq Universal SYBR Green Supermix (Bio-Rad) and primers obtained from the Fly Primer Bank listed in Table S1. The percent knockdown for each gene in question was derived from two individual parent sets and each of these biological replicates was run with two technical replicates on the CFX96 Real Time PCR Detection System (Bio-Rad) for a total of four samples per gene. RpL32 (RP-49) was used as reference genes and the knockdown efficiency (%) was obtained using the ΔΔCq calculation method compared to the control (*vermilion* RNAi only). Sep2^2^ and Sep5^2^ were previously shown to be loss-of-function mutants ^48,54^. Sep1 RNAi, Sep4 RNAi, Pnut RNAi, and Anillin RNAi knockdown efficiencies are 98%, 93%, 99%, and 94%, respectively.

### qPCR

Total RNA was obtained from 100 embryos (0-30 min old) using TRIzol (Invitrogen). 1 µg of total RNA was used for reverse transcription with the iScript™ gDNA Clear cDNA Synthesis Kit (Bio-Rad). RT-PCR analysis was performed using the iTaq™ Universal SYBR® Green Supermix (Bio-Rad) with two individual parent sets and two technical replicates on the CFX96TM Real Time PCR Detection System (Bio-Rad). RpL32 was used as a reference gene. The % knockdown was calculated using the ΔΔCq calculation method compared with control (vermilion knockdown). The same primer set for RpL32 was used in previously described ^80^.

### Embryo handling and preparation

Nuclear cycle 4-6 embryos were collected for 30min at 25°C and harvested at room temperature (22°C). Collected embryos were dechroionated by hand, mounted onto No. 1.5 coverslips coated with glue, and covered with Series 700 halocarbon oil (Halocarbon Products Corp.) as previously described ^12^.

### Laser wounding

All wounds were generated with a pulsed nitrogen N2 Micropoint laser (Andor Technology Ltd., Concord, MA, USA) tuned to 435 nm and focused on the cortical surface of the embryo. A region of interest was selected in the lateral midsection of the embryo and ablation was controlled by MetaMorph. On average, ablation time was less than 3s, and time-lapse imaging was initiated immediately. Occasionally, a faint grid pattern of fluorescent dots is visible at the center of wounds that arises from damage to the transparent vitelline membrane that covers embryos.

### Microscopy

All imaging was performed at room temperature (22°C). The following microscopes were used:

1. For live imaging: Revolution WD systems (Andor Technology Ltd., Concord, MA, USA) mounted on a Leica DMi8 (Leica Microsystems Inc., Buffalo Grove, IL, USA) with a 63x/1.4 NA objective lens and controlled by MetaMorph software. Images and videos were acquired with 488 nm and 561 nm, using an Andor iXon Ultra 897 or 888 EMCCD cameras (Andor Technology Ltd., Concord, MA, USA).
2. A Yokogawa CSU-X1 confocal spinning disc head mounted on a Nikon Eclipse Ti (Nikon Instruments, Melville NY,USA) with a 60x/1.4 NA objective lens and controlled by MetaMorph software. Images and videos were captured using 488nm and/or 561nm lasers with a Hamamatsu C9100-13 EMCCD camera. All images for cell wound repair were 17-20 µm stacks/0.25 µm steps. For single color, images were acquired every 30 sec for 15 min and then every 60 sec for 25 min or every 10 sec for 3 min. For dual green and red colors, images were acquired every 30 sec for 30-40 min. Live imaging for premature ooplasm streaming was performed as previously described ^82^.
3. For actin polymerization assays and fixed tissues: Zeiss LSM 780 spectral confocal microscope (Carl Zeiss Microscopy GmbH, Jena, Germany) fitted with Zeiss 20x/0.8 Plan-Apochromat, 40x/1.4, and 63×/1.4 oil Plan-Apochromat objectives. FITC (Alexa 488) fluorescence was excited with the 488 nm line of an Argon laser, and detection was between 498-560 nm. Red (Alexa 568) fluorescence was excited with the 561 nm line of a DPSS laser and detection was between 570-670 nm. Far-red (Phalloidin 633) fluorescence was excited with the 633 line of an Argon laser, and detection was between 570-670 nm. Confocal sections were acquired at 0.25-1.0 micron spacing.

### Image processing, analysis, and quantification

All images were analyzed with Fiji ^83^. Measurements of wound area were done manually. To generate xy kymographs, all time-lapse xy images were cropped to 5.8 µm x 94.9 µm and then each cropped image was lined up. Fluorescent line profiles were generated and graphed using Matlab (Mathworks) as previously described ^6^. The lines represent the averaged fluorescent intensity and gray area is the 95% confidence interval. Line profiles from the left to right correspond to the top to bottom of the images unless otherwise noted.

Line profiles showing protein recruitment relative to the actin ring were generated similar to previously described with some alterations. The Septin and actin line profiles across the wound at 120sec were first normalized to the respective reference unwounded profiles. The results were then averaged about the center of the wound resulting in circumferentially averaged profiles. Each set of profiles (actin and Septin) were then averaged (n≥10) and normalized to their respective maximum values and graphed with the gray area depicting 95% confidence interval and the center of the wound (left) and the outer edge of the wound (right) to show protein recruitment with respect to the actin ring.

Wound expansion was calculated with max wound area/initial wound size. Closure rate was calculated with two time points, one is t_max_ that is the time of reaching maximum wound area, the other is t<half that is the time of reaching 50-35% size of max wound since the slope of wound area curve changes after t<half. Using these time points, average speed was calculated with (wound area at t_max_ – wound area at t<half)/t_max_-t<half.

Quantification of the width and average intensity of actin ring was performed as follows: the width of actin ring was calculated with two measurements, the ferret diameters of the outer and inner edge of actin ring at 120 sec post-wounding. Using these measurements, the width of the actin ring was calculated with (outer ferret diameter – inner ferret diameter)/2. The average intensity of the actin ring was calculated with two measurements. Instead of measuring ferret diameters, we measured area and integrated intensity in the same regions as described in ring width. Using these measurements, the average intensity in the actin ring was calculated with (outer integrated intensity – inner integrated intensity)/(outer area – inner area). To calculate relative intensity for unwounded (UW) time point, average intensity at UW was measured with 507#x00D7;50 pixels at the center of the embryos and then the average intensity of the actin ring at each timepoint was divided by the average intensity of the UW measurement. To quantify protein recruitment to wounds, the fluorescent intensity of the UW image was subtracted from the image of the ring at 50% closure. We then generated the fluorescence profile of the subtracted image and calculated the area under the curve (AUC) using Matlab (Mathworks), similar to previously described ^80^. Generation of all graphs and Kruskal Wallis test were performed with Prism 8.0 (GraphPad Software Inc.) or Matlab (Mathworks).

### Protein expression

Sep1, Sep2, Pnut, and Sep5 cDNAs were amplified as 5′BamHI-3′NotI fragments from Drosophila cDNAs RE30523, LD20082, LD37170, and LD28935, respectively. Sep4 cDNA was amplified as 5’Sal1-3’Not1 fragment from LD37170. The cDNA clones come from the Drosophila Gene Collections 1 or 2. These all were cloned into a double-tag pGEX vector (GST and His) ^84^. Each Septin construct was transformed into Rosetta-gami B(DE3) cells. For Sep1-Sep2-Pnut complexes, Sep1 and Sep2 cDNAs were amplified and cloned into a pETDuet vector. For Sep4-Sep5-Pnut complexes, Sep4 and Sep5 cDNAs were amplified and cloned into a pETDuet vector. Pnut cDNA was amplified and cloned into a pET28-double-tag vector (GST and His). pETDuet-Septins and pET28-pnut constructs were transformed into BL21(DE3)pLysS cells. Primers used for cloning are described in Table S1.

Protein expression assays were performed as previously described ^85^. For Septin proteins, cells were lysed by sonication in lysis buffer (500mM NaCl, 5 mM MgCl_2_, 50 mM Tris pH 8, 40 μM GDP, and 1 mM DTT) with 0.5% Triton X-100, 1% Deoxycholate, 50 mM imidazole, and complete protease inhibitor tablets (Roche). Lysates were centrifuged at 10,000 *g* for 30 min, and the supernatants were coupled to Fastflow His-sepharose (GE) overnight at 4°C. The matrix was washed three times with lysis buffer with 50 mM imidazole and eluted by lysis buffer with 1.5 M imidazole. The His elutions were subsequently coupled to glutathione-sepharose 4B (GE) overnight at 4°C, then washed with lysis buffer and released from the beads by cutting with PreScission protease (GE) diluted in PreScission Protease Buffer (500 mM NaCl and 50 mM Tris HCl). All proteins were dialyzed into lysis buffer and then flash frozen.

Rho family GTPase proteins were expressed using a modified pGEX vector transformed into Rosetta-Gami B BL21(DE3) pLysS strains (Novagen). Protein was purified from these cells by freeze-thaw lysis in Hepes buffer (20mM Hepes; pH 7.5, 150mM NaCl, 5% glycerol) with Complete protease inhibitor tablets (Roche) or Phenylmethylsulfonyl fluoride (PMSF). Triton X-100 was added to 1% and lysates were sonicated as required. Lysates were centrifuged at 10,000 rpm for 30 minutes in a JA-17 or SS-34 rotor and the supernatants were coupled to Glutathione-sepharose 4B (GE) for 3 hours at 4°C. The matrix was washed three times with Hepes buffer then stored at 4°C.

### F-actin polymerization assays

Slides and coverslips were cleaned with water then 100% ethanol. The slides were then treated with a nitrocellulose solution (1:1 solution of sterile Collodion in 2% amyl acetate and amyl acetate; Electron Microscopy Sciences) and then left to airdry for one hour. Polymerization assays were performed using a mixture of commercial rhodamine labeled actin monomers (Cytoskeleton, Inc.) and unlabeled actin monomers that were mixed at a 1:1 ratio. This mixture was then diluted to 100mM concentration in G-Buffer (10 mM Tris pH7.5, 50 mM KCL, 2 mM MgCl_2_, 1 mM ATP). 0.5µl of this actin mixture was added to a tube that contained purified Septin proteins at the concentration of (100nM), 1 mM Trolox, 2 mM protocatechuic acid, 0.1 μM protocatechuate 3,4-dioxygenase and 0.1% (w/v) methylcellulose at a final volume of 10µl. After thorough mixing, 6µl of this solution was added to a slide and covered with a nitrocellulose treated coverslip and sealed. The slides were incubated at room temperature for an hour before imaging to ensure that the actin filaments had time to polymerize. The slides were visualized by confocal microscopy using a Zeiss LSM 780 at 40x or 60x with a 2x zoom. Fields of view were 40µm x 40µm.

### GST Pulldowns

GST pulldowns were performed similar to previously described ^85^. GST-proteins bound to glutathione-sepharose were washed with Hepes-LS buffer (20mM Hepes; pH 7.5, 150mM NaCl, 10% glycerol, 0.1% Triton X-100). Test proteins were generated *in vitro* using the TNR quick-coupled transcription-translation kit (Promega). For radiolabeling, ^35^S-Methionine was added to the IVT reaction. IVT lysates post-translation were diluted in Hepes-LS + Pepstatin, Aprotinin, Leupeptin, and AEBSF and pre-incubated with GST on glutathione Sepharose for one hour at 4°C to eliminate non-specific binding species. The pre-cleared lysates were then added to GST-protein in Hepes-LS and incubated for one hour at 4°C. The Sepharose matrix was then washed 5 times with Hepes-LS and the bound fraction analyzed by SDS-PAGE followed by autoradiography. In each case, 1% input (post clearing with GST) is shown.

GST-Rho family GTPase fusion proteins were exchanged while bound to Glutathione-sepharose by incubating with either GTP, GDP, or GDP-pNPP in exchange buffer (50mM HEPES pH 7.08, 20mM MgCl_2_ 5mM EDTA, 0.1mM EGTA, 50mM NaCl, 0.1mM DTT) for 30 minutes at 30°C. Exchange was performed immediately prior to use in pulldown assays.

### Statistical analysis

All statistical analysis was done using Prism 8 (GraphPad, San Diego, CA). Gene knockdowns were compared to the appropriate control, and statistical significance was calculated using a Kruskal Wallis test with *p*<0.05 considered significant unless otherwise noted.

---

**Fig S1.**
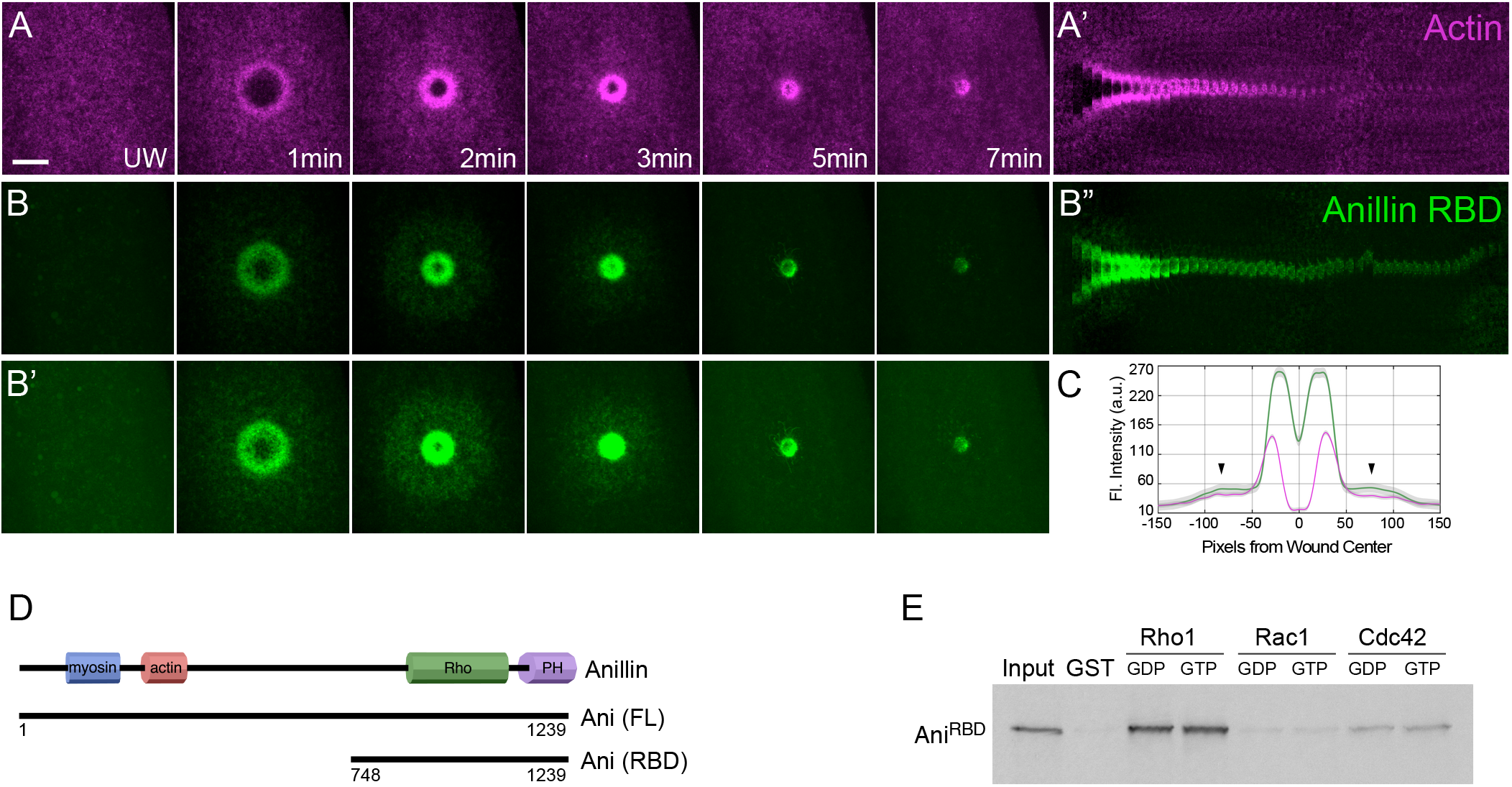
The Anillin RBD biosensor overlaps with both the actomyosin ring and actin halo, and interacts with Rho1 and Cdc42 in a nucleotide-independent manner. (A-B) Confocal max projection images of wounds generated in embryos expressing an actin reporter (sStMCA; A) and the Anillin RBD-GFP biosensor (B). (B’) Over-exposed view of the image shown in B. Dashed circle indicates recruitment of the Anillin RBD biosensor to the actin halo region. Scale bar: 20 µm. (C) Fluoresence intensity (arbitrary units) profile across the wound area in A-B. Arrowheads indicate the recruitment of the Anillin RBD biosensor to the actin halo region. (D) Schematic diagram of the Drosophila Anillin protein showing the extent of the full-length and RBD fragments used in these studies. (E) GST pull-down assays with 35S-labeled *in vitro* translated Anillin RBD protein with bacterially purified GST alone or GDP- or GTP-loaded Rho1, Rac1, or Cdc42 proteins as indicated. The Anillin RBD fragment binds to Rho1, and to a lesser extent Cdc42, in a nucleotide-independent manner.

## VIDEO LEGENDS

**Video 1.** Septins exhibit distinct localization patterns in cell wound repair. Time-lapse confocal xy images and fluorescence intensity (arbitrary units) profiles across the wound area from *Drosophila* NC4-6 staged embryos co-expressing an actin reporter (sStMCA or cGMCA, shown in magenta) along with Sep1-GFP (green) (A), Sep2-GFP (green) (B), ChFP-Pnut (shown in green) (C), Sep4-GFP (green) (D), or Sep5-RFP (shown in green) (E). Time post-wounding is indicated. UW: unwounded.

**Video 2.** Septin knockdowns/mutants exhibit distinct phenotypes in cell wound repair. (A-G) Time-lapse confocal xy images and fluorescence intensity (arbitrary units) profiles across the wound area from *Drosophila* NC4-6 staged embryos expressing an actin marker (sGMCA) in: control (*vermilion* RNAi; A), Sep1 RNAi (B), *Sep2*^2^ mutant (C), Pnut RNAi (D), Sep4 RNAi (E), *Sep5*^2^ mutant (F), or Sep4 RNAi + Pnut RNAi (G). Time post-wounding is indicated. UW: unwounded.

**Video 3.** Anillin is recruited to wounds and necessary for cell wound repair. (A) Time-lapse confocal xy images and fluorescence intensity (arbitrary units) profiles across the wound area from *Drosophila* NC4-6 staged embryos co-expressing an actin reporter (sStMCA, magenta) along with GFP-Anillin (green). (B) Time-lapse confocal xy images and fluorescence intensity (arbitrary units) profiles across the wound area from *Drosophila* NC4-6 staged embryos expressing an actin marker (sGMCA) in Anillin RNAi. (C) Time-lapse confocal xy images and fluorescence intensity (arbitrary units) profiles across the wound area from *Drosophila* NC4-6 staged embryos co-expressing an actin reporter (sStMCA, magenta) along with Anillin^RBD^-GFP biosensor (green). Time post-wounding is indicated. UW: unwounded.

## Key resources table

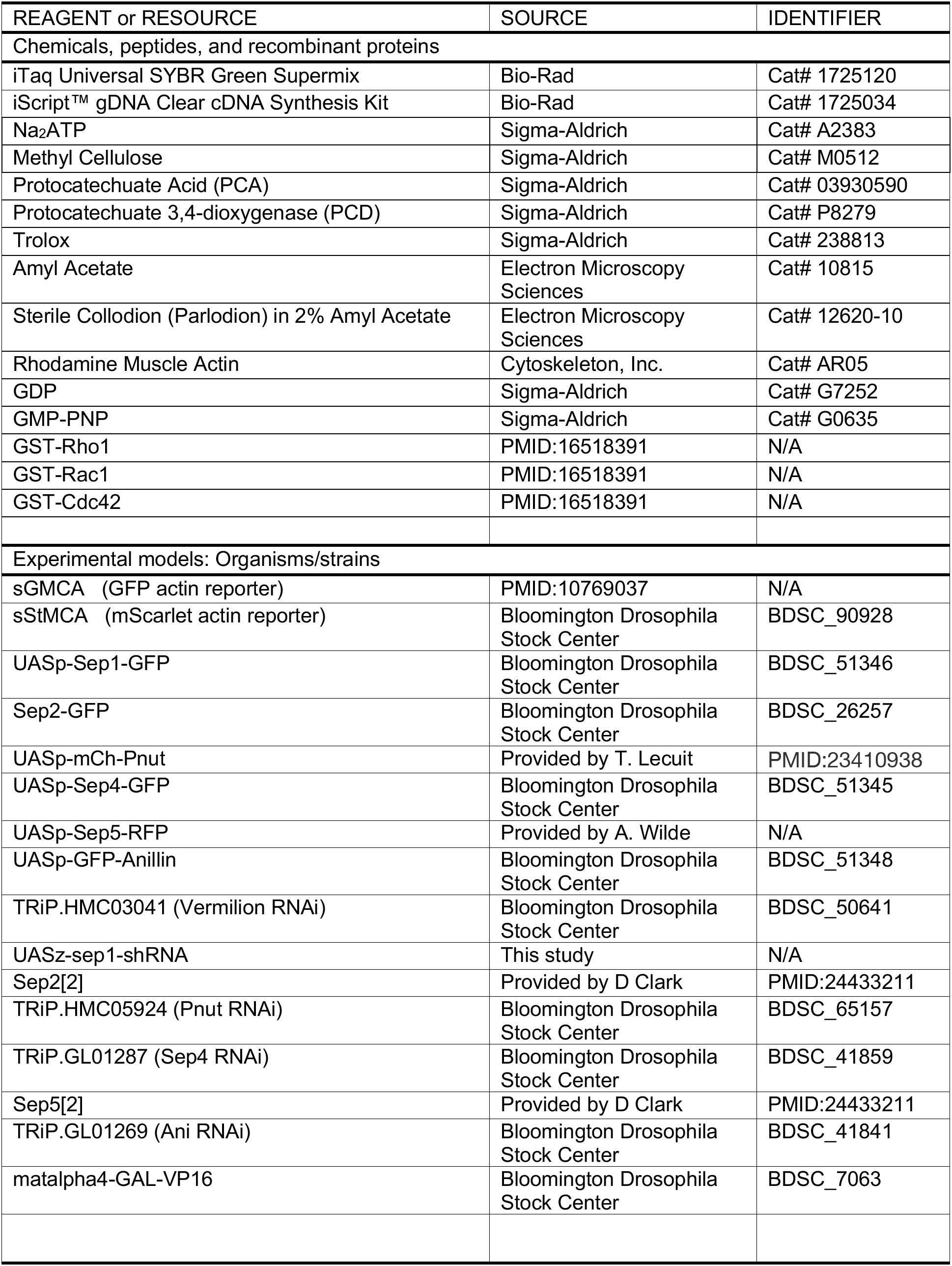
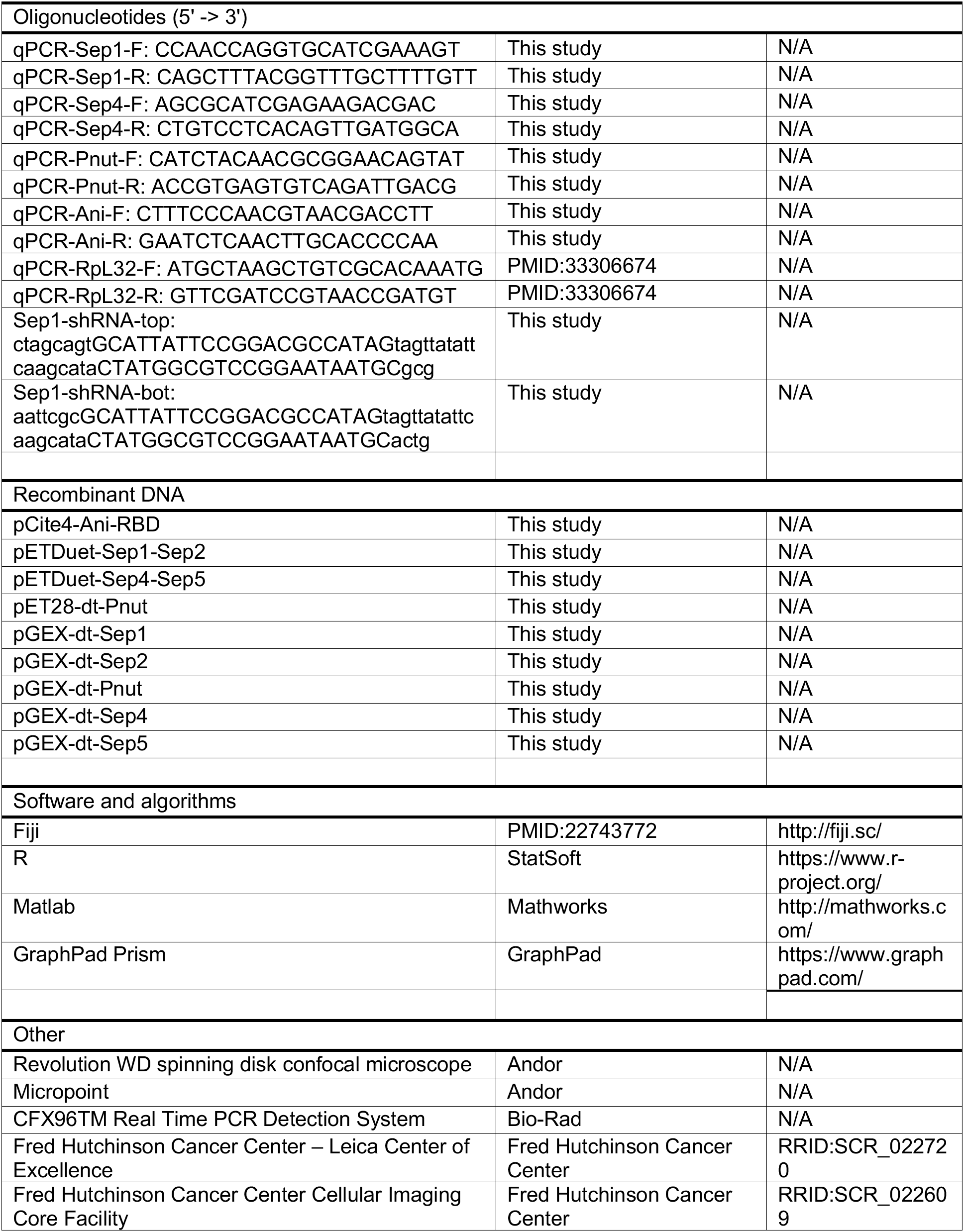

